# Opto-Myomatrix: μLED Integrated Microelectrode Arrays for Optogenetic Activation and Electrical Recording in Muscle Tissue

**DOI:** 10.1101/2024.07.01.601601

**Authors:** Jiaao Lu, Muneeb Zia, Danish A. Baig, Young Jin Lee, Euichul Chung, Jeong Jun Kim, Geyu Yan, Kailash Nagapudi, Philip Anschutz, Shane Oh, Daniel H. O’Connor, Samuel J. Sober, Muhannad S. Bakir

## Abstract

**Objective:** Optogenetics is a valuable and widely-used technique that allows precise perturbations of selected groups of cells with high temporal and spatial resolution by using optical systems and genetic engineering technologies. This study aims to develop Opto-Myomatrix, a novel optogenetic tool for precise muscle fiber control and high-resolution electrophysiological recording.

**Method:** Based on a flexible and biocompatible polymer substrate, the device incorporates an integrated µLED that delivers light at 465 nm for optogenetic stimulation and 32 PEDOT:PSS-coated electrodes for electromyography (EMG) recording. We also added a reflector to improve optical power output.

**Results:** The fabricated Opto-Myomatrix device achieves an optical output intensity as high as 129.46 mW/mm^2^ in the direction of interest, which is enhanced by nearly 100%. The PEDOT:PSS-coated electrodes exhibit 85% lower impedance than uncoated contacts, enabling high signal-to-noise EMG acquisition. We investigated heat dissipation characteristics of the µLED through measurements and finite element simulations, confirming that temperature changes remain within safe limits. The Opto-Myomatrix device was implanted in transgenetic mice and successfully stimulated targeted jaw muscles, inducing movement while simultaneously capturing EMG signals.

**Conclusion:** The Opto-Myomatrix effectively combines precise optical stimulation and high-quality EMG recording in a flexible and biocompatible device, focusing on optogenetic applications in muscle.

**Significance:** While optogenetic tools are well-established for brain and central nervous system (CNS) research, the development of Opto-Myomatrix addresses a critical gap by enabling precise muscle fiber control and high-resolution recording for advancing neuromuscular studies.

## I. Introduction

Optogenetics is a unique technique that incorporates genetic engineering and optical systems to control the activity of cells with high precision. This technique employs opsins, which are light-sensitive proteins that can be engineered for expression in cell membranes. These genetically modified cells can then be excited or inhibited using visible light at select wavelengths[1], [2]. One of the key advantages of optogenetics is its ability to precisely control neuronal spiking on a millisecond timescale [3], which is achieved by the rapid response of opsins to light. Such temporal resolution aligns closely with the natural timing of neural activity and is invaluable for studying dynamic neural processes. Additionally, optogenetics can target specific cell types within neural circuits by using cell-type-specific promoters, which direct opsin expression exclusively in designated cells [4]. This enables selective activation of the targeted muscle or nerve cells, avoiding less precise control seen with other neuralmodulation techniques such as deep brain stimulation (DBS) or functional electrical stimulation (FES). Another valuable feature provided by optogenetics is that several opsins can be expressed in a single cell, which allows selective excitation and inhibition at their corresponding wavelengths [5]–[7] This versatile manipulation of cellular activity further enhances the complexity and detail of cellular behavior studies.

Building on these foundational advantages, applying optogenetic techniques in muscle opens new avenues for understanding muscle function and disorders. The precision and specificity of optogenetics provide distinctive tools for mapping and controlling muscle fibers, thus allowing targeted treatments and rehabilitation strategies [8]. Most current research on optical stimulation in muscle fiber utilize lasers as a light source and separate fine-wire or needle electrodes to record electromyography (EMG) signals [9]–[13] However, although lasers have high light intensity and low beam divergence, they suffer from high power consumption, bulky size, and tethered optical and communication systems restricting the natural behavior of freely moving animals [1]. Moreover, finewire and needle electrodes primarily capture bulk EMG from multiple motor units, while for comprehensive computational analysis, it is crucial to identify signals from single motor units [14], [15]. Hence, in muscle optogenetics, there is a need for devices capable of simultaneous light delivery and electrophysiological recording to investigate the causal relationship between neuronal activation and behavior. A promising alternative light source is µLEDs, which are often chosen in optical neural implants developed for brain tissues, due to their compact size, lightweight design, low power consumption, and rapid modulation capability [16]–[For electrical signal acquisition from muscle, high-density, flexible microelectrode arrays (MEA) are a recently developed tool positioned directly on muscle fiber that can detect motor units activated by a single motor neuron. Incorporating µLEDs into flexible MEA provides the means to obtain accurate recording of neural activities corresponding to optical stimulation with different spatial distributions. It also alleviates the complexity of the surgery process as only one device needs to be implanted[21].

In this work, we propose Opto-Myomatrix, a µLED-integrated flexible microelectrode array, to realize simultaneous optical stimulation in muscle fiber and high-resolution EMG recording. The device consists of 32-channel recording electrodes and a µLED chip to deliver light stimulation at 465 nm. In order to increase optical intensity in the direction of interest, an aluminum reflector underneath the µLED chip was fabricated, and optical characterization of the device was conducted, demonstrating enhanced optical output power suitable for optogenetics stimulation in skeletal muscle. To further improve electrochemical performance, poly(3,4-ethylenedioxythiophene)-poly(styrenesulfonate) (PEDOT:PSS) was applied on all recording electrodes to achieve lower impedance, hence achieving a high signal-to-noise ratio. Both thermal characterization and finite element simulation of the µLED on a flexible array were presented to verify the temperature rise from the device remains safe for muscle stimulation. Finally, we implanted the Opto-Myomatrix device in transgenic mice to optically stimulate jaw muscles, inducing jaw movement, and to collect EMG signals at the same time.

## II. Design and fabrication

### A. Design concept

Fig.1 presents the schematic of the Opto-Myomatrix: one µLED chip for optical stimulation is positioned at the center of an array of 32 electrodes for EMG recording. The µLED chip (CREE^®^ TR2227™, Cree, Inc., USA) has a peak wavelength at 465 nm, which falls within the activation range for Channelrhodopsin-2 (ChR2)[3]. It is connected to two traces for power supply and encapsulated by transparent polydimethylsiloxane (PDMS). Given that the bare µLED chip emits light in every direction, we introduce a reflector placed underneath the µLED (as shown in the blue inset in Fig.1). This design aims to redirect the light emitted from the backside forward, thereby enhancing the optical power output in the desired direction. The 32 recording electrodes surrounding the µLED each has a footprint of 200*×*100 µm^2^ and are coated with a thin layer of PEDOT:PSS (yellow inset in Fig.1) to improve electrochemical performance. All the metal traces and electrodes are sandwiched between two layers of flexible and biocompatible polyimide. The thicknesses of the substrate and the encapsulation layers are 10 µm and 5 µm, respectively. A shallow trench is etched on the top polyimide layer to place the µLED. The size of the trench is designed to appropriately fit the chip with a footprint of 220*×*270 µm^2^ and a thickness of 50 µm.

**Fig. 1.**
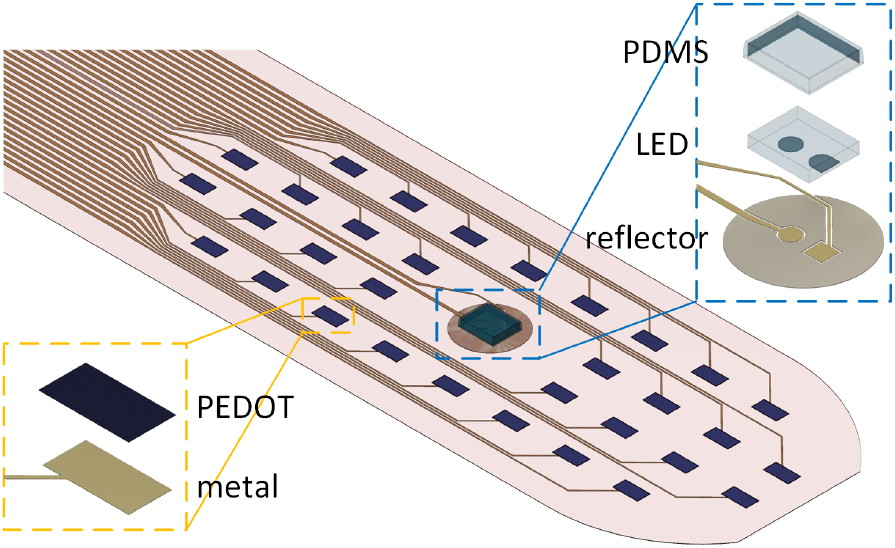
3D schematic of the Opto-Myomatrix. The device consists of 32 recording electrodes coated with PEDOT:PSS and a µLED chip at the center.

### B. Fabrication process

The fabrication process for Opto-Myomatrix is adapted from previous works on flexible MEAs[22]–[24] and is summarized in Fig 2. A detailed fabrication procedure is described in the following sections.

#### 1) Flexible microelectrode array

A 4-inch silicon wafer was utilized as the carrier substrate. After cleaning by acetone, methanol and isopropanol consecutively, and drying by compressed nitrogen, the wafer was spin-coated with a layer of polyimide (Pyralin 2611, HD MicroSystems™ L.L.C., USA) (Fig. 2(a)). The spin speed was set at 1500 RPM to precisely control the thickness of the bottom layer to be 10 µm. This specific thickness ensures the polymer substrate maintains both flexibility and durability through out the *in vivo* implantation process. The polyimide was then cured in a programmable oven at a maximum temperature of 320°C for 45 minutes. To form the traces and electrodes on the polyimide layer, negative photoresist (AZ® nLOF 2020, Microchemicals GmbH, Germany) was patterned, followed by thermal evaporation of titanium (20 nm) and gold (300 nm), and lift-off of the films in acetone (Fig. 2(b)). Subsequently, another layer of polyimide was formed by spinning at 4000 RPM and curing in an oven, serving as the top insulation layer. The top polyimide layer was etched with a reactive ion etching (RIE) process using a photoresist mask (MICROPOSIT™ S1827®, Kayaku Advanced Materials, Inc., USA), to expose the recording electrodes, contact pads for µLED, and connector pads (Fig. 2(d)). Next, an aqueous solution of PEDOT:PSS (PH 1000 PEDOT:PSS, Oscilla, UK) was spin-coated on the wafer and patterned via a lift-off process (Fig. 2(e)). This step selectively coats PEDOT:PSS only on recording electrodes to achieve lower impedance.

**Fig. 2.**
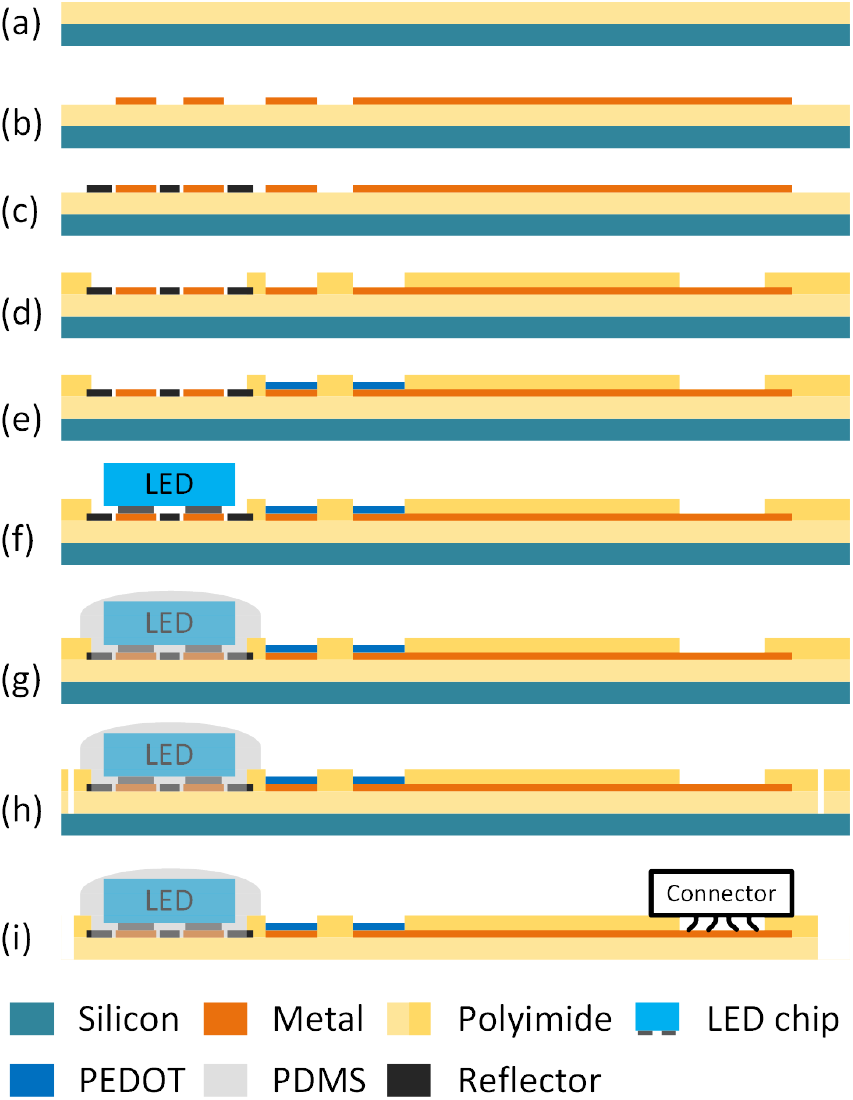
Fabrication process of Opto-Myomatrix. (a) Bottom polyimide coating. (b) Trace and electrode metal deposition. (c) Reflector metal deposition. (d) Top polyimide coating and patterning. (e) PEDOT:PSS coating and patterning. (f) µLED flip-chip bonding. (g) PDMS encapsulation. (h) Laser dicing. (i) Device release and connectorization.

#### 2) Reflection layer

As stated in Section II-A, the µLED chips have a colorless and transparent SiC substrate and emit light from all directions. However, in skeletal muscle stimulation, it is critical to maximize optical power output normal to the same face as the recording electrodes. Thus, we added a “mirror” layer underneath the LED chip to reflect back the light emitted towards the substrate of the device. The reflection layer consists of 100 nm of aluminum that was deposited and patterned via a lift-off process (Fig. 2(c)). A gap of 5µm was designed between the reflector and metal trace to avoid a short circuit path through aluminum. Reflectors with different shapes and sizes were fabricated and tested to investigate the most efficient geometry.

#### 3) µLED chip bonding and encapsulation

During the RIE process of the top polyimide layer (Fig. 2 (d)), shallow trenches of 230*×*280 µm^2^ were formed at the center of 32 electrode array. This process also exposed the contact pads to connect to the anode and cathode of µLED chip. The µLED chip was then attached to the wafer using a flip-chip bonder (FINEPLACER^®^ Lambda 2, Finetech GmbH & Co. KG, Germany) (Fig.2(f)). Thermocompression bonding was conducted by heating the chip to 290°C for 5 minutes and applying 8 N of force uniformly at the same time. Consequently, atomic interactions between the gold surface of the traces on the wafer and the anode/cathode on the µLED chip created metallic bonds at the interface, forming a strong and durable ohmic contact.

To encapsulate the µLED chips and to enhance the mechanical stability of the device, we carefully applied a thin layer of optically transparent PDMS only to cover the µLED chips (Fig.2(g)). A mixture of pre-polymer base and curing agent (SYLGARD™ 184 Silicone Elastomer Kit, Dow Inc., USA) was prepared at a ratio of 1:10. A small volume of the mixture was then dispensed onto the surface of µLED chip using a 30-gauge blunt-tip needle. The drop was allowed to spread to uniformly cover the µLED before placing the wafer in an oven for curing at 65°C for 2 hours under vacuum. This step is vital in preventing corrosion of the chips or deformation of the bonding due to moisture from tissue fluids when implanted *in vivo*.

#### 4) Device release and assembly

After µLED bonding, the outline of device was precisely cut through the polyimide layer using a femtosecond laser (OPTEC WS-Flex USP, OPTEC S.A., Belgium). Finally, the array was peeled off the silicon wafer to complete the release. A 32-channel Omnetics connector was assembled with anisotropic conductive film (ACF) tape for multichannel neural recording, while the µLED power was supplied by wires connected by conductive epoxy (Fig.2(i)).

### C. Fabricated device

Fig.3(a) and (c) show microscope and SEM images of the fabricated Opto-Myomatrix. The µLED chip is successfully flip-chip bonded on flexible MEA in the shallow trench on polyimide layer. When activated by current (Fig.3(a)), the µLED emits blue light at 465 nm, aligning with the activation wavelength for ChR2, thereby enabling targeted optogenetic stimulation. The blue inset in Fig.3(a) presents the circular reflector with diameter of 500 µm that is added underneath the µLED chip to enhance optical intensity of the Opto-Myomatrix device. The SEM image also shows that the top surface of the PDMS layer is above the surface of the polyimide substrate while the PEDOT:PSS layer of electrodes is slightly below the surface. This is due to the geometry of the µLED chip we selected and the etching process of top polyimide layer to expose recording electrodes. The surface profile of µLED chip covered by PDMS is captured using surface profilometry (Fig.3(b)). The maximum height at the center of the µLED chip is less than 60 µm. Considering that the thickness of the µLED chip is 50 µm, this maximum height indicates that the µLED chip is completely encapsulated by the PDMS layer, which has a thickness of less than 10 µm. The total thickness has been validated through *in vivo* experiments, ensuring that all electrodes can establish contact with muscle fibers and effectively record EMG signals (see Section IV). The released Opto-Myomatrix is shown in Fig. 3 (d), where the curve of the subsrate was formed by applying a bending force manually, demonstrating the device’s flexibility. As shown in the figure, the µLED is turned on with no loss in optical power. This indicates that the bending stress results in very little or no piezoelectric field change [25] and does not affect the bonding strength of the µLED chip, exhibiting a great potential for minimally invasive optical stimulation and physiological recording.

**Fig. 3.**
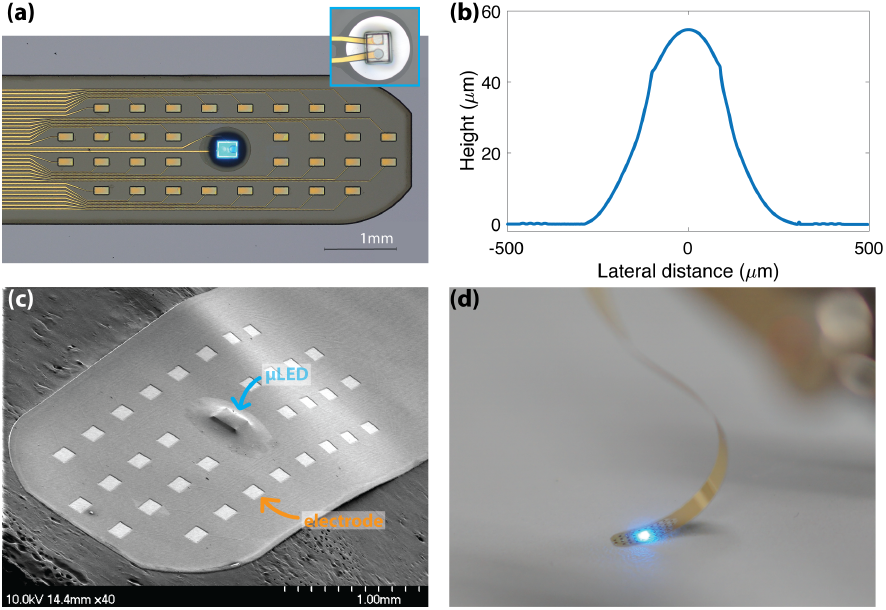
Fabricated Opto-Myomatrix. (a) A microscope image of all recording electrodes and µLED of Opto-Myomatrix on wafer; the inset shows the circular reflector with diameter of 500 µm. (b) Surface profilometry results for the µLED chip covered with a thin layer of PDMS on a flexible substrate. (c) An SEM image of the electrodes and µLED chip encapsulated in PDMS. (d) An optical image of released Opto-Myomatrix device under bending stress.

## III. Results

### A. Optical characterization

In order to evaluate the optical performance of fabricated Opto-Myomatrix and to assess the improvement from reflector design, optical power output from the µLEDs was measured at a wavelength of 465 nm with a power meter (PM100A, Thorlabs, Inc., USA) and a photodiode sensor (S120VC, Thorlabs, Inc., USA). Three types of reflectors were tested: circular shape with diameter of 500 and 1000 µm, and square shape with side length of 1000 µm. The µLED encapsulated in PDMS was placed as close as possible to the surface of the sensor, while a controlled DC power supply (E3610A 30-Watts DC Power Supply, HP Development Company, L.P., USA) was connected to supply current to the anode and cathode of the µLED. This setup aims to replicate the effect of stimulating muscle tissue, with the Opto-Myomatrix directly in contact with the muscle. The experiment setup was placed in air at room temperature and the optical intensity was calculated by dividing the measured power by total surface area of µLED (220*×*270 µm^2^).

As shown in Fig.4(a), the Opto-Myomatrix with bare µLED can achieve optical intensity of 5.93 to 66.20 mW/mm^2^ with supplied current from 1 to 20 mA. This indicates that at very low driving currents, our device exceeds the optical intensity typically required to activate ChR2 (1 mW/mm^2^)[26]. Additionally, the reflector design further increased the optical intensity. With the smallest reflector (500 µm dia.), the optical power is enhanced by at least 1.5x. Moreover, the larger circular reflector (1000 µm in diameter) increased the directed optical power by approximately 2x, with the highest intensity of 129.46 mW/mm^2^. Although the square reflector with 1000 µm side length has a larger footprint area, the average optical power measured is slightly lower than that with circular design.

**Fig. 4.**
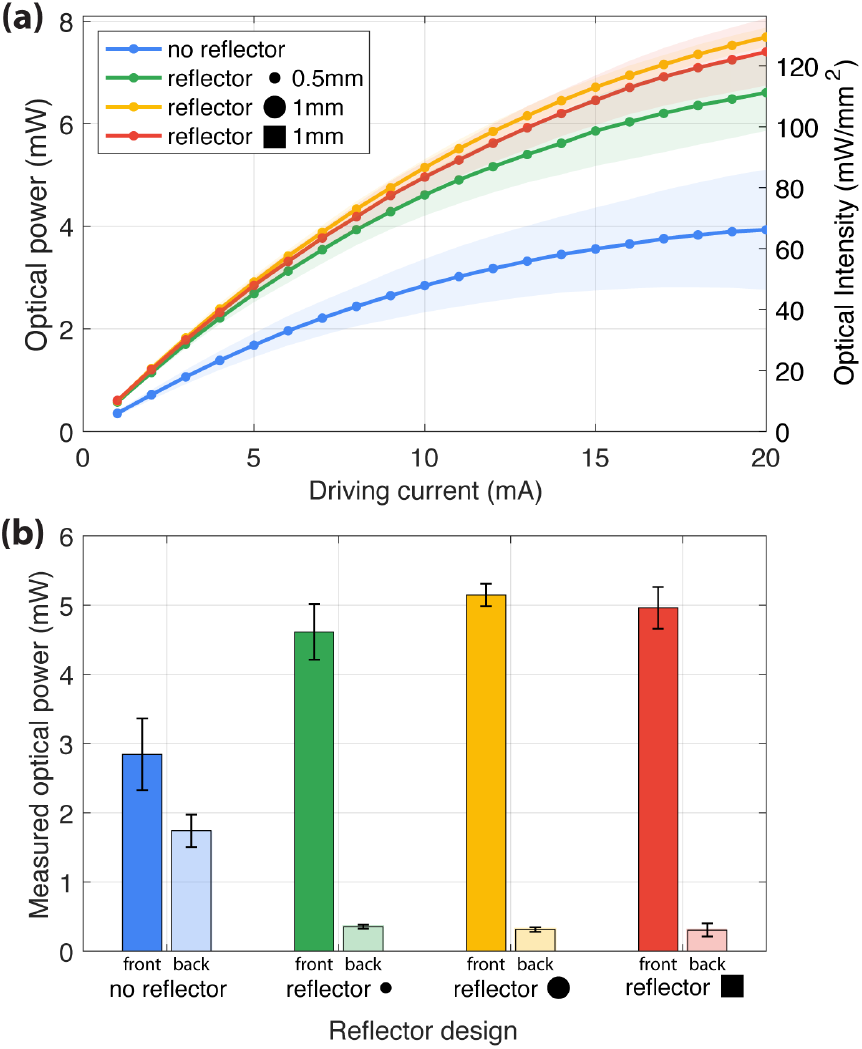
Optical measurement results. (a) Driving current vs. optical power output at 465 nm from the µLED with three types of reflectors (n=4 devices). (b) Optical power output measured from front and back side at 10 mA of supplied current with and without reflectors (n=4 devices).

As stated in Section II-A, the µLED chip employed in this work emits light in all directions. Hence, we conducted another set of measurements to obtain the optical power “leakage” from the backside of the device to further ensure the reflector design effectively prevents light leakage. In Fig.4 (b), constant current of 10 mA was supplied to the µLED with three different reflector designs, and the optical power output was read from the photodiode sensor placed towards the frontside and backside respectively. We also measured a µLED without reflector as the baseline result. The 1000 µm circular reflector gives the best frontside optical intensity (~2x baseline), and all reflector designs significantly reduce the optical power leaking from the backside (~5x baseline). Interestingly, the variation in the size of the reflector does not significantly affect its improvement on backside leakage, indicating that all reflectors are effective in redirecting light and enhancing the light intensity towards the targeted tissue.

### B. Electrochemical characterization

Electrochemical properties of the fabricated Opto-Myomatrix was evaluated by electrochemical impedance spectroscopy (EIS). Specifically, the recording electrodes with a bare gold surface and with PEDOT:PSS coating were tested, respectively, to evaluate the improvement in impedance from the conductive polymer layer. NanoZ impedance tester (White Matter LLC, USA) was employed to measure impedance from all 32 channels over the frequency range from 1 Hz to 5 kHz. The impedance data was obtained with a 2-electrode setup in physiological saline solution (0.9% NaCl), where the recording electrodes of Opto-Myomatrix acted as a working electrode (WE) and an Ag/AgCl wire served as reference electrode (RE) and counter electrode (CE).

Fig. 5 provides the EIS results measured from electrodes with both gold and PEDOT:PSS finishing. In line with our previous work [23], PEDOT:PSS has once again demonstrated its significant utility. Over the whole testing frequency range, PEDOT-coated electrodes consistently outperform the gold ones. Specifically, at 1 kHz, PEDOT-coated electrodes exhibit an average impedance as low as 3.4 kΩ, which represents an approximately 85% reduction compared with bare gold electrodes. This significant reduction in impedance suggests that PEDOT-coated electrodes can achieve a lower noise level, resulting in a higher signal-to-noise ratio in electrophysiology recordings [24]. Since the µLEDs were attached to the Opto-Myomatrix device after coating PEDOT:PSS layer, the measurement also confirms that the heat from the flip-chip bonding process does not compromise the integrity of the PEDOT:PSS layer.

**Fig. 5.**
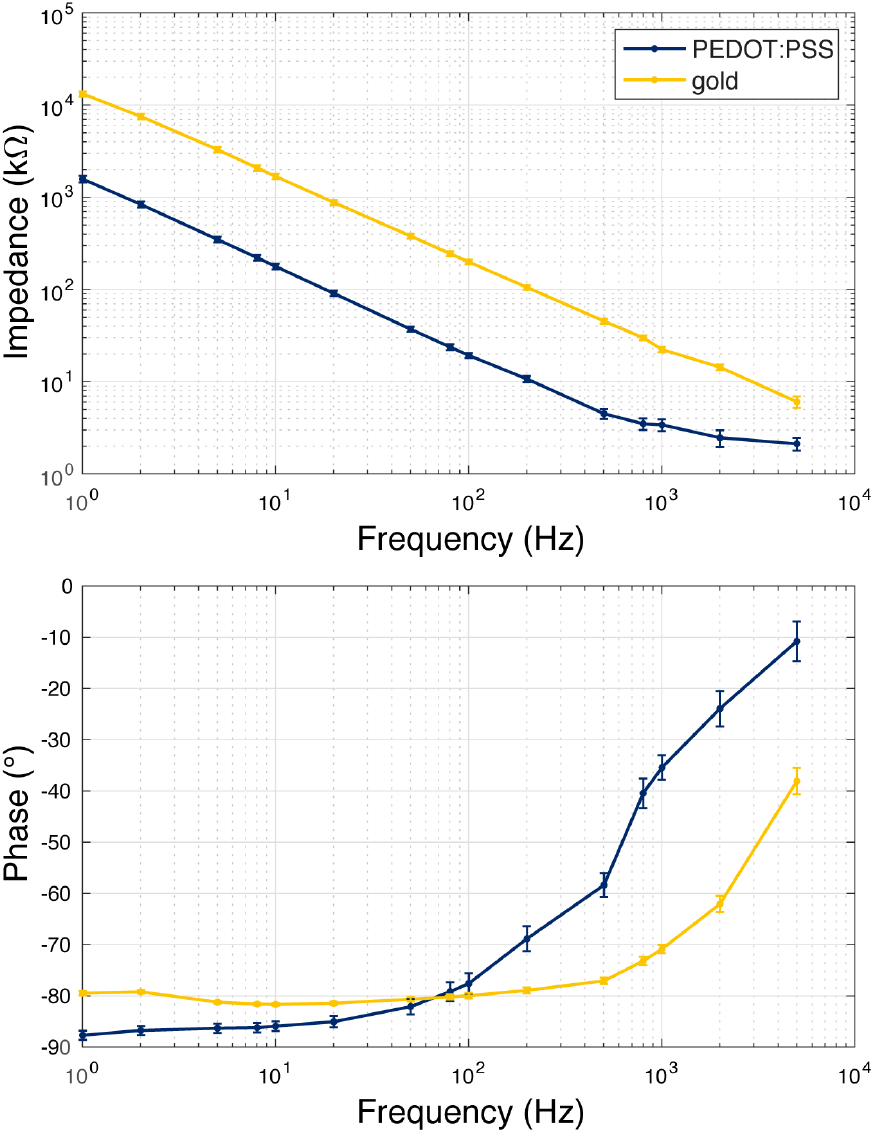
Electrochemical impedance spectroscopy (EIS) results of the recording electrodes on Opto-Myomatrix. Both bare gold and PEDOT:PSS coated electrodes were measured (n=32 electrodes).

### C. Thermal measurement

Temperature management is a critical consideration in muscle optogenetics stimulation to avoid adverse effects in muscle tissue [8]. The temperature profile of a single µLED on polyimide substrate was obtained using a thermal camera (A6701 MWIR, Teledyne FLIR LLC, USA) with a resolution of 640*×*512 and pixel size of 15 µm. A 1x microscope lens (3-5 µm, f/2.5 MWIR FPO Manual Bayonet Lens, Teledyne FLIR LLC, USA) was utilized and the system has a sensitivity of less than 0.02°C. This experiment was conducted in a controlled setting at room temperature (20°C). High-speed videos at 300 frames per second were recorded when a square-wave pulse current was supplied to the µLED with a duration of 20 ms and varying amplitude from 1 mA to 5 mA. To study the impact of reflector design on thermal performance, we conducted tests with three different device configurations: a bare µLED on the polyimide substrate, and µLEDs with two different sizes of circular reflectors (500 and 1000 µm in diameter). All µLEDs used in the tests were covered with a PDMS layer and the temperature variation was obtained from the surface of PDMS. To ensure accurate measurement of temperature, the samples were uniformly coated with high-emissivity black spray (emissivity, *ϵ* ≈ 0.95).

Fig. 6 (experiment) shows measured maximum temperature increase from different designs as a function of supplied current. For all device configurations, as the supplied current increases, more heat is generated, and thus the higher the temperature increase. By comparing the maximum temperature of the device with and without the aluminum reflector, it is noticed that the reflector acts as a heat spreader which effectively dissipates heat from the µLED, thereby reducing the temperature of the µLED. Furthermore, the device with a 1-mm diameter reflector exhibited a greater temperature reduction than the one with a 0.5-mm diameter reflector, as the spreading thermal resistance from µLED to reflector is inversely proportional to the diameter ratio of the reflector to the µLED (*R*_th,spread_ *~ D*_µLED_*/D*_reflector_) [27]. As a result, the device with a 1-mm reflector consistently maintains lower temperatures throughout the entire supplied current range. At supplied current of 5 mA, both reflectors successfully spread out the heat and obtained a controlled temperature increase of 2.08°C and 2.68°C, respectively, while the maximum temperature increase from bare µLEDs covered with PDMS reached 4.14°C. Although previous research indicates that a temperature increase of 2°C can potentially damage brain tissue [17], [28], [29], muscle tissue has a higher tolerance for temperature fluctuations compared to neural tissue, allowing it to withstand up to 3°C without sustaining damage [30], [31]. In this scenario, only using the highest current (5 mA) without a reflector surpasses the safe limit. It is important to mention that the measurements here were conducted in air, which has a much lower thermal conductivity than muscle tissue [32]. Consequently, when the Opto-Myomatrix is implanted *in vivo*, the surrounding muscle tissue is expected to more effectively dissipate the heat generated by the µLED chip. This will likely allow for the safe operation of the Opto-Myomatrix with a higher supplied current than used in these experiments.

**Fig. 6.**
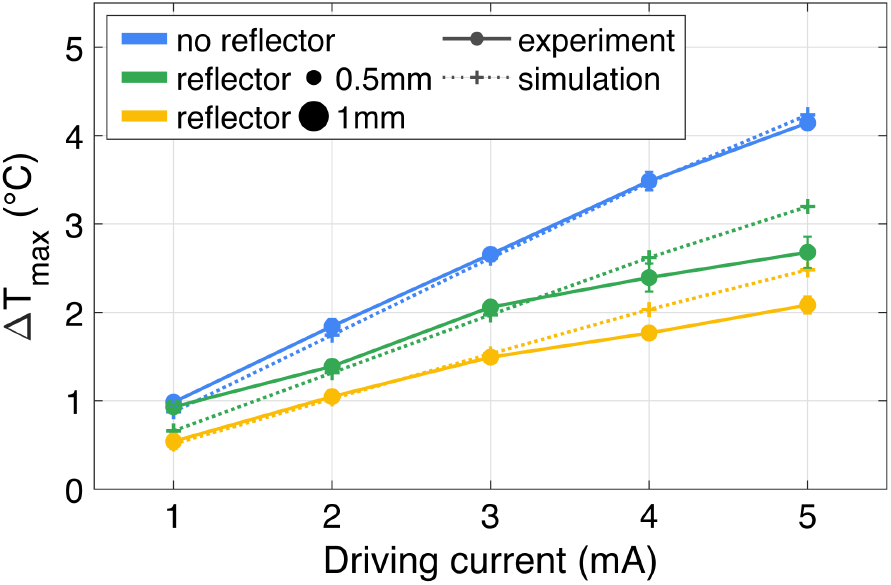
Temperature measurement of µLED chip operating with 20 ms current pulse at different amplitudes (n=3 devices), along with finite element simulation results of the experiment.

A thermal finite element analysis (FEA) was also performed using ANSYS® Mechanical APDL 2024 R2 to model the experimental setup for the measurements. As shown in Fig.6, numerical simulation results align well with the experimental data, with the maximum error of less than 0.5°C. The discrepancies primarily arise from geometric simplifications in the numerical model and possible inaccuracies in estimating the actual heat dissipation from the µLED.

### D. Thermal finite element analysis

Since it is challenging to experimentally obtain the temperature profile within the muscle when the opto-Myomatrix is implanted *in vivo*, thermal finite element analysis of the opto-Myomatrix within muscle tissue was carried out. To reduce computing complexity, the model is simplified to contain the main components: the µLED chip made of silicon carbide, the anode and cathode of µLED made of gold with a size of 80 µm diameter, the polyimide substrate with a size of 1.1*×*1.1 mm^2^, the PDMS covering µLED chip with a thickness of 10 µm, and a substantial volume of muscle tissue with a volume of 5×5×5 cm^3^. Fig.7(a) shows the simplified geometry used for finite element analysis, illustrating half of the device cut along the symmetry plane at *x* = 0. The physical properties of the materials used in the simulated models are detailed in Table I (sourced from the ANSYS® material library). The PDMS and polyimide surfaces are in contact with muscle to replicate *in vivo* experiment setup. The muscle boundaries are set at constant body temperature (37°C) except for surfaces in contact with air, assuming the volume for analysis is adequately large. All the other boundaries that are exposed to air are configured for natural convection in air, with a heat transfer coefficient of 5 W/(m^2^ K) at room temperature (20°C)[33]. The heat source is applied at the bottom side of the µLED chip to mimic the heat generated at the p-n junctions, and the input power is estimated from the I-V curve of the chip assuming 40% optical efficiency (*Q* = *I·V* × (1 −40%))[34]. We selectively simulated square-wave current pulse with a duration of 5, 10, and 15 ms and amplitude from 1 to 5 mA, and studied the transient response to extract the highest temperature increase at the surface of muscle in contact with the µLED. Similar to the thermal measurement experiment, the reflector is taken into account as well to investigate its potential function as a heat spreader.

**Fig. 7.**
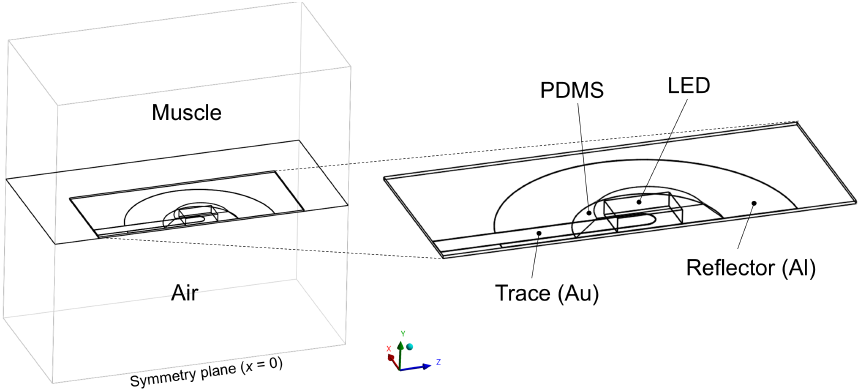
Model geometry used in the thermal finite element analysis.

**TABLE 1.** PHYSICAL PROPERTIES OF TISSUE AND ENGINEERING MATERIALS USED IN THE SIMULATIONS.

|  | Density<br>(kg/m<sup>3</sup>) | Heat capacity<br>(J/kg·K) | Thermal conductivity<br>(W/m·K) |
| --- | --- | --- | --- |
| Silicon carbide | 3200 | 1200 | 450 |
| Gold | 19300 | 129 | 317 |
| Polyimide | 1300 | 1100 | 0.15 |
| PDMS | 970 | 1460 | 0.16 |
| Aluminum | 2700 | 904 | 237 |
| Muscle | 1090 | 3421 | 0.49 |

**TABLE 2.** COMPARISON OF KEY PROPERTIES IN SELECTED OPTOGENETIC NEURAL IMPLANTS.

|  |  | Cao *et al.*<br>[21] | Wu *et al.*<br>[19] | Scharf *et al.*<br>[39] | Ji *et al.* [18] | Ayub *et al.*<br>[40] | Taal *et al.*<br>[41] | **This work** |
| --- | --- | --- | --- | --- | --- | --- | --- | --- |
| Optical | LED type | LED chip | Customized GaN μLED | Customized GaN μLED | μLED chip from Cree | Customized GaN μLED | Organic LED | μLED chip from Cree |
|  | Wavelength (nm) | 465 | 460 | 450 | 470 | 455 | Blue/orange | 465 |
| | Emission area ($\mu\text{m}^2$) | $6 \times 10^5$ | 150 | 491 | $5.9 \times 10^4$ | 1964/7854 | 399 | $5.9 \times 10^4$ |
| | Intensity ($\text{mW}/\text{mm}^2$) | 0.7 | 2.8 | 400 | 241.7 | 2.23 | 0.1/0.25 | 129.5 |
|  | # of channels | 1 | 12 | 64 | 16 | 10 | 1024 | 1 |
| | Driving current | 5 mA | 15-40 $\mu\text{A}$ | 0.2-1.1 mA | 1-20 mA | 0.04-2 mA | 35 $\mu\text{A}$ | 1-20 mA |
| Electrical | Contact material | Au | Ir | - | IrOx | - | PEDOT:PSS | PEDOT:PSS |
| | Size of electrode ($\mu\text{m}^2$) | 5000 | 143 | - | $3.14 \times 10^4$ | - | 1024 | $2 \times 10^4$ |
|  | # of channels | 3 | 32 | - | 16 | - | 32 | 32 |
| | Impedance@1kHz (kΩ) | - | 1000 | - | 3 | - | - | 3.4 |
| General | Thermal analysis | - | FEA | Thermal imaging | Thermal imaging | Integrated thermistor | - | Thermal imaging and FEA |
|  | Substrate material | Polyimide | Silicon | Silicon | Polyimide | Silicon | Silicon | Polyimide |
|  | Application | Cortex | Hippocampus | Cortex | - | Cortex | Cortex | Muscle fiber |

Fig.8 shows the overall thermal characteristics of the opto-Myomatrix implanted within muscle tissue. Fig.8(a) presents the maximum temperature difference (Δ*T*_max_) for each device as a function of applied current for different pulse durations, where Δ*T*_max_ is defined as the maximum temperature at the muscle-contacting surface minus body temperature (37°C). As expected, the maximum temperature increases with longer pulse durations and higher currents, due to increased heat accumulation within the µLED. Notably, all reflector configurations successfully keep the maximum temperature increase below 2°C, even with the longest pulse duration (15 ms) and highest current (5 mA). This demonstrates the heat-spreading performance of the reflector design. All devices, regardless of the presence of a reflector, exhibit temperature variations within 3°C under various current and pulse durations, indicating a safe operating range for muscle optogenetics stimulation [30], [31]. Fig. 8(b) presents the transient thermal response of the devices at a fixed current of 3 mA and pulse duration of 15 ms. The reflectors effectively lower the peak temperature and accelerate the temperature decline following the completion of the 15 ms current pulse, indicating improved heat-spreading capability. Fig. 8(c) shows the transient temperature variation for the device with reflector D = 1 mm at a current of 3 mA, when the pulse duration increases from 5 to 10, 15, and 20 ms. Longer pulse durations result in higher peak temperatures and extend the cooling time required to return to ambient temperature.

**Fig. 8.**
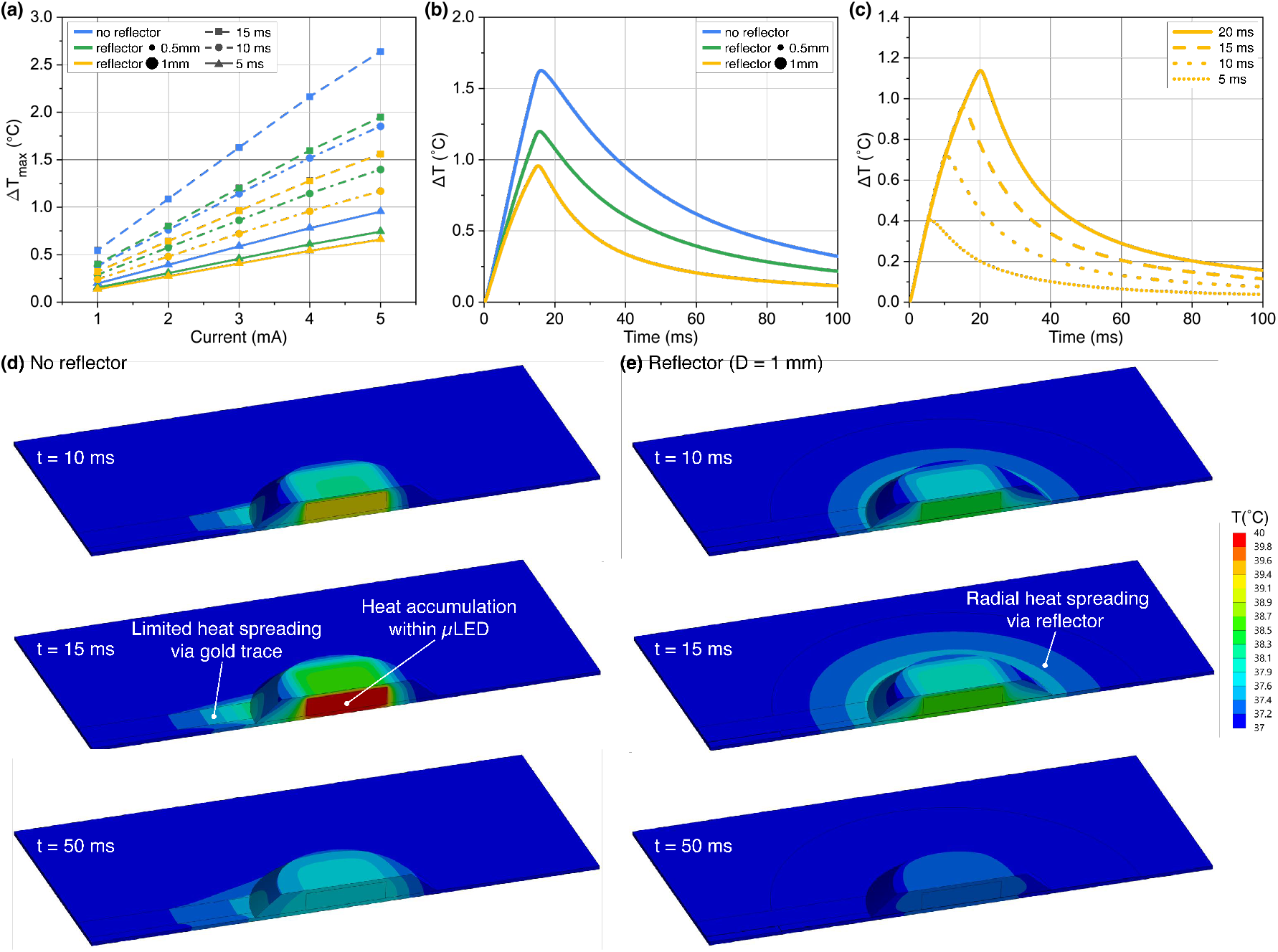
Thermal finite element analysis of the µLED on an Opto-Myomatrix device. (a) Maximum temperature increase simulated at various current amplitudes and pulse durations. (b) Transient thermal response of the devices when the µLED is supplied with current of 3 mA for 15 ms. (c) Transient thermal response of Reflector (D = 1 mm) device with current of 3 mA for various pulse durations. (d, e) Temperature distributions of the device without reflector and with reflector (D = 1 mm) at different time points; current amplitude: 3 mA, pulse duration: 15 ms.

To further investigate the cooling mechanism of the aluminum reflector, the temperature distribution within the device and at surfaces in contact with muscle tissue was analyzed over time. Figs. 8(d) and (e) show the temporal temperature distribution of devices without and with a reflector (denoted as ‘No reflector’ and ‘Reflector (D = 1 mm)’), under a current of 3 mA and a pulse duration of 15 ms. Temperature contours at 10 ms, 15 ms, and 50 ms are presented to visualize temperature rise and subsequent decrease. We omit the contours in the surrounding muscle and air regions to focus on the temperature at the interface between muscle tissue and the device. Comparing the temperature contours reveals that the 1-mm reflector facilitates radial heat spreading, maintaining lower temperatures relative to the no-reflector case. Without a reflector, heat conduction is limited primarily to the gold traces on the substrate, leading to substantial heat accumulation and higher temperatures at the µLED. Given that aluminum has a thermal conductivity (denoted as *k*) over three orders of magnitude higher than that of polyimide (*k*_Al_*/k*_PI_ = 1580), the reflector creates an effective radial conduction path that significantly reduces temperatures at the interface with muscle tissue[35]. Fig.9 shows the temperature distribution within the muscle tissue, with the cross-sectional and top views for the device with reflector (D = 1 mm) at 15 ms under the same conditions.

**Fig. 9.**
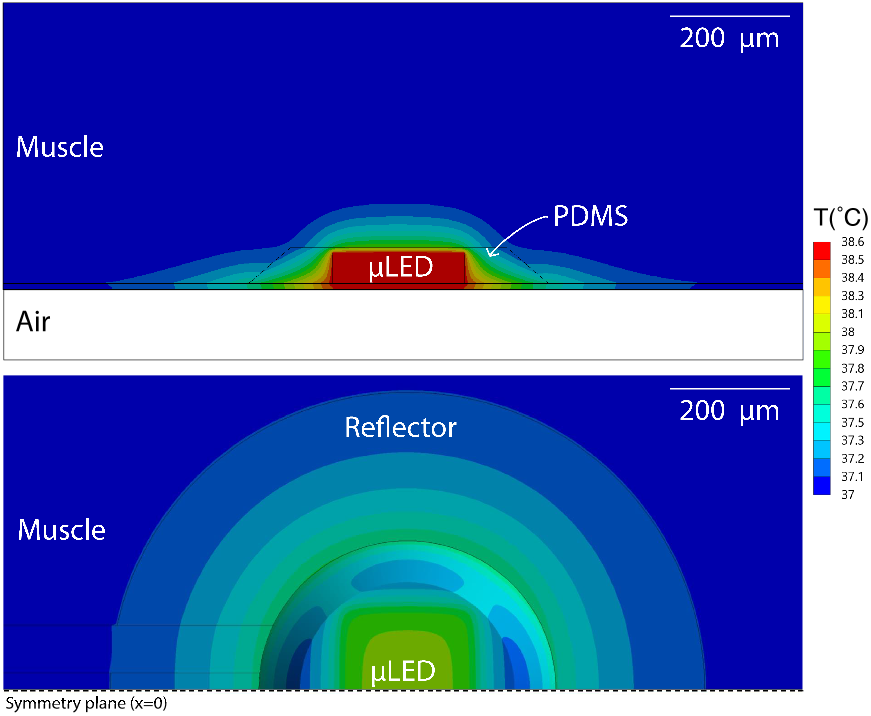
Cross-sectional and top views of temperature distribution in the device with reflector (D = 1 mm); current amplitude: 3 mA, pulse duration: 15 ms.

## IV. *In vivo* experiments

The fabricated Opto-Myomatrix device was utilized in an acute electrophysiological recording in mice to optically stimulate temporalis muscle and record EMG signals (Fig.10(a)). All procedures were approved by the Johns Hopkins University Institutional Animal Care and Use Committee.

Emx1-Cre/Ai32 offspring were generated by crossing Emx1-IRES-Cre knock-in homozygotes (Stock no. 005628, Jackson Laboratory) with Ai32 (RCL-ChR2H134R-EYFP) reporter homozygotes (Stock no. 024109, Jackson Laboratory). The mice were anesthetized using 0.9-1.5% isoflurane at a flow rate of 1 L/min and the anesthesia was maintained consistently throughout the surgical procedure and data collection. 30 µL of physiological saline was injected subcutaneously, followed by opening a skin flap at the chin to expose the temporalis and masseter muscles. The Opto-Myomatrix devices were then placed on the surface of the temporalis muscles while the temporalis muscle was kept moist with saline. A highspeed CMOS camera (DR1-D1312-200-G2-8, PhotonFocus, Switzerland) in a 0.25x telecentric lens (Edmund Optics, USA) captured jaw kinematics at 400 frames per second using IR illumination at 850 nm and an angled mirror to collect side and bottom views simultaneously. Light pulses of varying power (0-1 mW) and pulse duration (5, 50, 100 ms) were delivered via the integrated µLED component. Stimuli parameters were controlled by a National Instruments DAQ system (BNC-2090A, National Instruments Corp., USA) with WaveSurfer. EMG signals were amplified and digitized with an Intan RHD2216 bipolar amplifier chip (Intan Technologies, USA) and bandpass filtered at 300–7500Hz.

The collected raw EMG signals were rectified and then smoothened with a Gaussian kernel of 2.5 ms. To quantify the optically evoked jaw movement, eleven keypoints were traced along the jaw in captured video using DeepLabCut to quantify the movement [37]. The illumination of ChR2-expressing temporalis muscles, a jaw closing muscle, leads to elevation and deviation of the mandible away from the midline [38]. As shown in Fig 10(b), short (5 ms) light pulses of *<*1 mW power is sufficient to drive overt jaw movements. Increasing light power drives increasing amplitude in jaw movements. Correspondingly, the higher light power also results in higher peak EMG amplitude and lower latency in EMG activity, suggesting greater activation of muscle fibers and decreased time to depolarization.

**Fig. 10.**
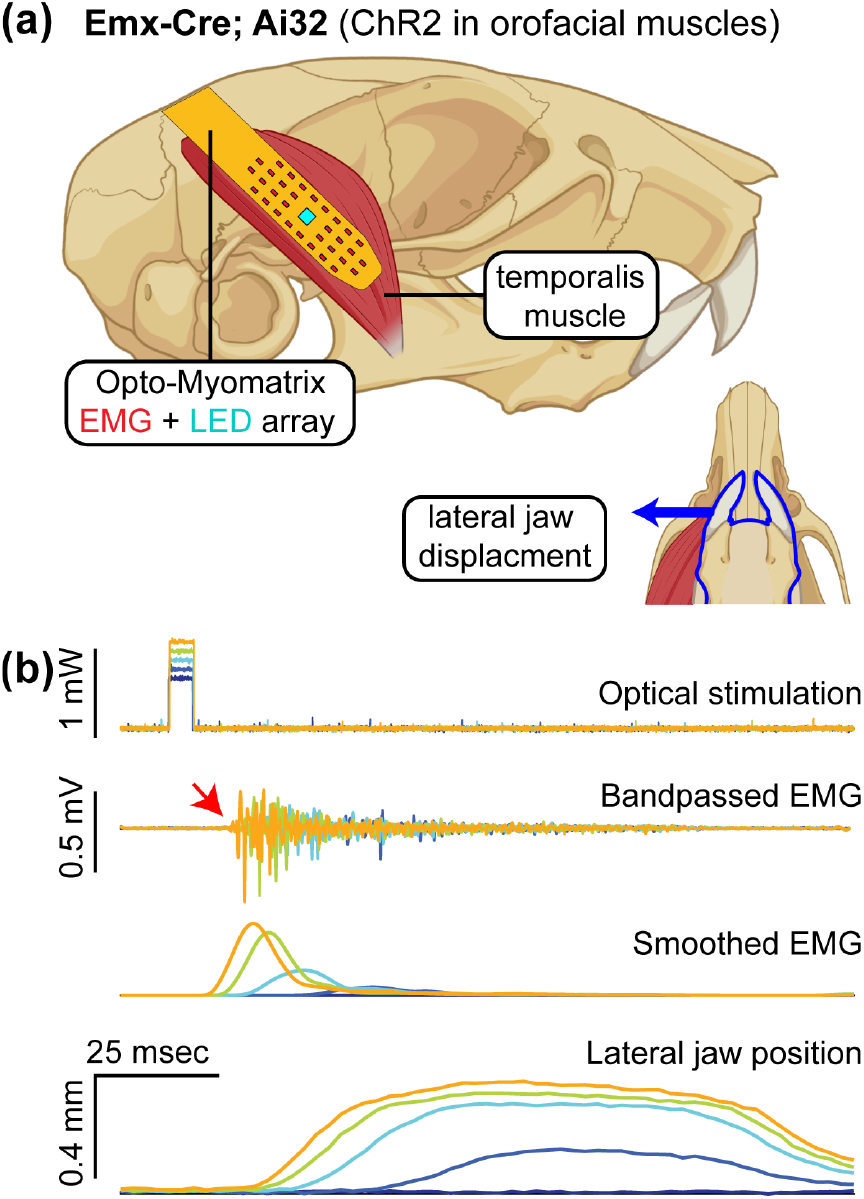
*In vivo* optical stimulation. (a) Experiment setup. Optical stimulation of the temporalis muscle in an anesthetized mouse expressing channelrhodopsin-2 (ChR2) in orofacial muscle fibers. (b) Optical activation evokes muscle activation and lateral movement of the jaw. The latency of EMG response (red arrow) is consistent with published values for optical stimulation of muscle fibers[36].

The multi-contact layout of Opto-Myomatrix (Fig. 11(a)) permits measurement of the spatial extent of optical depolarization of muscle. The space constant was derived via a single-term exponential fit using the equation *y* = *ae*^*−x/λ*^, where *y* represents the peak EMG amplitude, *x* the distance from the *µ*LED, and *λ* the space constant. As shown in Fig. 11(b), the peak amplitude of EMG activity decayed with distance from centrally placed µLED with a space constant of approximately 1111 um. The space constant reflects the spread of depolarization across muscle fibers. The spatial profile of EMG activity was notably inhomogeneous, likely reflecting the underlying orientation of muscle fibers and variability in electrode contact with the underlying muscle. In addition, the spatial profile of latency in EMG activity showed increasing latency with distance from µLED.

**Fig. 11.**
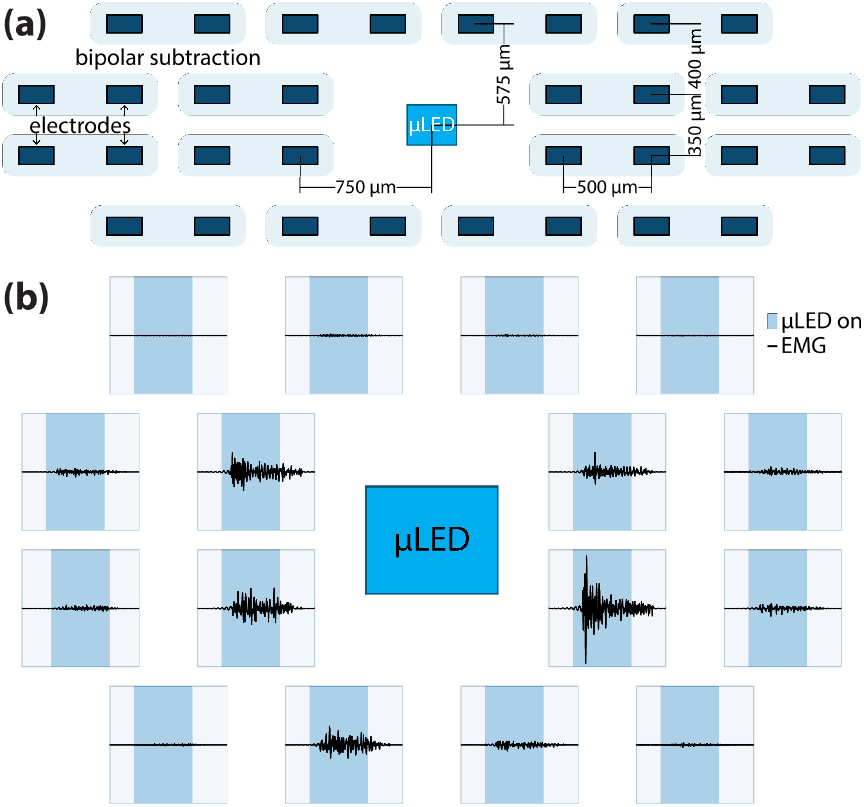
Sample EMG signals. (a) Layout of 32-channel recording electrodes paired for bipolar recording and µLED on Opto-Myomatrix. (b) EMG signals recorded using the Opto-Myomatrix in bipolar mode during optical stimulation. The pulse is 50 msec in duration and each plot spans from −0.5 to +0.5 mV. Variations in amplitude and latency across the channels are illustrated, highlighting their correlation on the varying distances from the µLED.

## V. Discussion

We compared the key properties of optical, electrical, and overall performance of Opto-Myomatrix with several state-of-the-art neural implants using LED technology for optogenetic applications (see Table 2). With a commercially available chip µLED, our device achieves a relatively high optical intensity while maintaining a low-complexity and cost-effective fabrication process. It features a high channel count and low electrode impedance, which are advantageous for acquiring stable, low-noise signals in muscle recordings. The thermal performance was investigated with both thermal imaging and finite element analysis (FEA) to validate that the device can be used safely during stimulation of muscle. While the other devices primarily target cortical or hippocampal regions, we presents a novel approach by focusing on muscle tissue, broadening the scope of optogenetics beyond traditional neural studies. To our knowledge, this is the first flexible device with integrated light source designed specifically for optical activation of muscle fibers.

However, we have only validated the Opto-Myomatrix with acute *in vivo* experiment lasting less than one day. In the future, we plan to conduct long-term performance evaluations to assess the device’s durability in chronic applications. Specifically, we will focus on improving hermetic sealing techniques with PDMS and other material such as parylene-C, to enhance the device’s longevity and stability in prolonged use.

## VI. Conclusion

Opto-Myomatrix is a flexible and biocompatible microelectrode array with an integrated µLED chip for simultaneous optogenetic stimulation and EMG recording from muscle tissue. In this work, we present the design concept, fabrication process and characterization of Opto-Myomatrix, and proof-of-concept *in vivo* experiments to validate its capability of precisely controlling and monitoring muscle activity. The fabrication process can be adapted to fit various muscle types, enhancing the versatility of the device. Featuring an embedded reflector design, the Opto-Myomatrix achieves an optical intensity of up to 129.46 mW/mm^2^ at wavelength of 465 nm with a 20 mA driving current, significantly surpassing the activation threshold for ChR2. Additionally, the reflector plays a crucial role in minimizing light leakage from the backside, redirecting the light to the targeted area. The electrochemical characterization shows that the PEDOT:PSS-coated recording electrodes of Opto-Myomatrix possess low impedance, consistently outperforming the benchmark device with gold finishing over the testing frequency range. The observed impedance improvement aligns with the consistent findings of reduced impedance demonstrated in our previous work [23], enabling recordings with a high signal-to-noise ratio and thereby enhancing both data quality and reliability. We also conducted thermal measurements and simulated the heat dissipation process of the µLED chip and confirmed that the maximum temperature increase stays within safe limits for muscle stimulation.

The fabricated Opto-Myomatrix was implanted in transgenic mice and successfully stimulated temporalis muscle with light at 465 nm. The induced EMG signals and jaw movement were collected and analyzed to demonstrate the device’s functionality and the precision of optogenetic muscle control. The multicontact layout of Opto-Myomatrix also allowed quantification of optical depolarization across muscle fibers, showing a space constant of 1111 um and a distance-related variation in EMG activity and latency.

## Acknowledgment

This work was funded by National Institutes of Health Grant R01 NS109237 and U24 NS126936, the McKnight Foundation and the Simons Foundation as part of the Simons-Emory International Consortium on Motor Control, and was performed in part at the Georgia Tech Institute for Electronics and Nanotechnology (IEN) and Materials Characterization Facility (MCF); JJK was supported by NIH Award 1F31DE033256-01.

## Notes

### Competing Interest Statement

The authors have declared no competing interest.

### Summary of Updates

Section III C and D updated on thermal measurement and simulation; Section V added.

